# Detection of cytosolic *Shigella flexneri* via a C-terminal triple-arginine motif of GBP1 inhibits actin-based motility

**DOI:** 10.1101/212175

**Authors:** Anthony S. Piro, Dulcemaria Hernandez, Sarah Luoma, Eric. M. Feeley, Ryan Finethy, Azeb Yirga, Eva M. Frickel, Cammie F. Lesser, Jörn Coers

## Abstract

Dynamin-like guanylate binding proteins (GBPs) are gamma interferon (IFNγ)-inducible host defense proteins that can associate with cytosol-invading bacterial pathogens. Mouse GBPs promote the lytic destruction of targeted bacteria in the host cell cytosol but the antimicrobial function of human GBPs and the mechanism by which these proteins associate with cytosolic bacteria are poorly understood. Here, we demonstrate that human GBP1 is unique amongst the seven human GBP paralogs in its ability to associate with at least two cytosolic Gram-negative bacteria, *Burkholderia thailandensis* and *Shigella flexneri.* Rough lipopolysaccharide (LPS) mutants of *S. flexneri* co-localize with GBP1 less frequently than wildtype *S. flexneri*, suggesting that host recognition of O-antigen promotes GBP1 targeting to Gram-negative bacteria. The targeting of GBP1 to cytosolic bacteria, via a unique triple-arginine motif present in its C-terminus, promotes the co-recruitment of four additional GBP paralogs (GBP2, GBP3, GBP4 and GBP6). GBP1-decorated *Shigella* replicate but fail to form actin tails leading to their intracellular aggregation. Consequentially, wildtype but not the triple-arginine GBP1 mutant restricts *S. flexneri* cell-to-cell spread. Furthermore, human-adapted *S. flexneri,* through the action of one its secreted effectors, IpaH9.8, is more resistant to GBP1 targeting than the non-human-adapted bacillus *B. thailandensis*. These studies reveal that human GBP1 uniquely functions as an intracellular ‘glue trap’ inhibiting the cytosolic movement of normally actin-propelled Gram-negative bacteria. In response to this powerful human defense program *S. flexneri* has evolved an effective counter-defense to restrict GBP1 recruitment.

**Importance:** Several pathogenic bacterial species evolved to invade, reside and replicate inside the cytosol of their host cells. One adaptation common to most cytosolic bacterial pathogens is the ability to co-opt the host’s actin polymerization machinery, in order to generate force for intracellular movement. This actin-based motility enables Gram-negative bacteria such as *Shigella* to propel themselves into neighboring cells thereby spreading from host cell to host cell without exiting the intracellular environment. Here, we show that the human protein GBP1 acts as a cytosolic ‘glue trap’ capturing cytosolic Gram-negative bacteria through a unique protein motif and preventing disseminated infections in cell culture models. To escape from this GBP1-mediated host defense, *Shigella* employs a virulence factor that prevents or dislodges the association of GBP1 with cytosolic bacteria. Thus, therapeutic strategies to restore GBP1 binding to *Shigella* may lead to novel treatment options for shigellosis in the future.

## Introduction

Cell-autonomous immunity describes the ability of a single cell to defend itself against intracellular pathogens and constitutes an essential branch of the immune system (1, 2). Cell-autonomous immunity in vertebrates is often orchestrated by IFN-inducible genes (ISGs) (2). Amongst the most robustly expressed ISGs are those encoding dynamin-like guanylate binding proteins (GBPs) (3–5). GBPs control intrinsic antiviral, antiprotozoan and antibacterial immunity, are highly expressed in inflamed tissue, and can be predictive of infectious disease progressions (5–10). Since their discovery, seven human *GBP* orthologs and one pseudogene have been identified. The *GBP* genes are located within one gene cluster on chromosome 1 (11). Other vertebrate genomes contain comparable numbers of *GBP* orthologs: *e.g.* mice possess 11 genes in addition to 2 pseudogenes (12). Human and mouse GBPs share a high degree of homology with the most conserved region found within their N-terminal G domains. However, GBP protein family members are highly divergent from each other at their very C-terminal ends, both within and across different vertebrate species (11). The functional consequence of this C-terminal amino acid sequence variability has not been previously explored.

To exert many of their anti-microbial functions, GBPs specifically associate with intracellular microbes residing in the host cell cytosol or at pathogen-occupied supramolecular structures, which include viral replication complexes (10) and pathogen-containing vacuoles (3–5). Following pathogen recognition, GBPs are thought to deliver antimicrobial host factors to pathogen-containing vacuoles and to bacteria residing in the host cell cytosol, thereby enabling the execution of distinct defense pathways which include membranolytic destruction of cytosolic bacteria (13, 14), capture of microbes within degradative autolysosomes (7), and activation of inflammasomes (13–19). These studies clearly demonstrated the importance of GBPs in host defense against a broad spectrum of pathogens in a vertebrate host. However, because these previous studies were conducted almost exclusively in mouse models, it remains to be determined whether human GBPs execute functions comparable to their murine counterparts.

In this study we systematically tested all seven members of the human GBP protein family for their ability to co-localize with the cytosolic bacterial pathogens *Listeria monocytogenes, Shigella flexneri* and *Burkholderia thailandensis.* All of these bacterial species are equipped with the ability to co-opt the host actin polymerization machinery for actin-based cytosolic motility and cell-to-cell spread (20). While we failed to detect any co-localization between human GBPs and the Gram-positive bacterium, *L. monocytogenes,* we found that GBP1, independent of other human GBP paralogs, targeted both Gram-negative bacteria, *B. thailandensis* and *S. flexneri*. This specific interaction between GBP1 and bacteria, which is determined by a unique C-terminal triple-arginine motif, inhibits actin-tail formation of GBP1-decorated bacteria resulting in a reduction of bacterial cell-to-cell spread. We further observed that GBP1 targeted the non-human-adapted microbe *B. thailandensis* more efficiently than the human-adapted pathogen *S. flexneri* and identified the bacterial ubiquitin E3 ligase IpaH9.8 as a *S. flexneri* virulence factor that interferes with GBP1 recruitment to this professional cytosolic bacterium. Thus, this study provides a novel understanding of the role of human GBPs in immunity to bacterial pathogens and also defines a virulence strategy employed by a human-adapted microbe to escape from GBP1-regulated host defense.

## Results

### Human GBP1 co-localizes with cytosolic *S. flexneri* and *B. thailandensis*

Most known GBP-mediated antimicrobial functions require that GBPs directly localize to intracellular microbes or their surrounding vacuoles (3–5). Based on this premise we screened the entire set of human GBPs (GBP1 – 7) as ectopically expressed mCherry N-terminal fusion proteins in the human epithelial lung carcinoma cell line A549 for co-localization with the cytosolic bacterial pathogen *S. flexneri.* Expression of each fusion protein was detectable by Western blotting (Figure S1) and by immunofluorescence (Figure 1A). Unexpectedly, only a single member, GBP1, associated with cytosolic *S. flexneri,* as assessed at 1 and 3 hours post infection (hpi) in either naïve or IFNγ-primed A549 cells (Figure 1A). Association of GBP1 with individual *S. flexneri* bacteria was observed as early as 10 minutes post host cell invasion (data not shown and video S1). Similarly, we observed that GBP1 was the sole human GBP family member to co-localize with *B. thailandensis,* another cytosolic Gram-negative pathogen (Figure 1B). In contrast to the observed targeting of GBP1 to these Gram-negative bacteria, we failed to detect any co-localization with *L. monocytogenes,* a cytosolic Gram-positive pathogen (Figure 1C).

**Figure 1.**
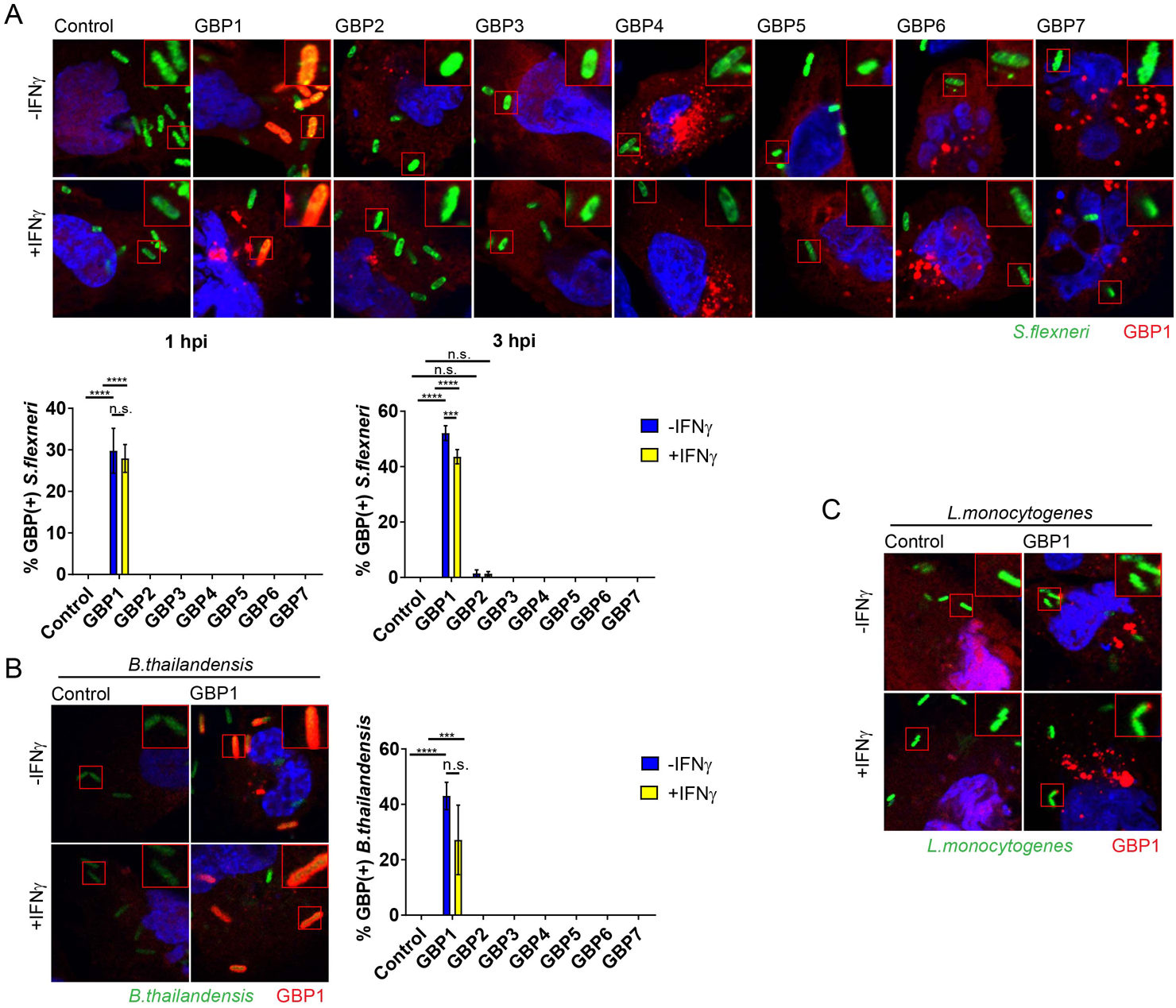
Ectopically expressed human GBP1 co-localizes with Gram-negatives *S. flexneri* and *B. thailandensis* but not Gram-positive *L. monocytogenes* in human A549 cells. A549 cells were transfected with indicated GBP paralogs fused to mCherry at their N-termini or transfected with mCherry control. Cells were primed with IFNγ at 200 U/ ml overnight or left untreated. (A) Cells were infected with GFP-positive *S. flexneri* at a multiplicity of infection (MOI) of 50 and microscopy images were taken at 1 hpi. (B) Cells were infected with GFP-positive *B. thailandensis* at an MOI of 100 and images were acquired at 8 hpi. (C) Cells were infected with GFP^+^ *L. monocytogenes* at an MOI of 5 and monitored at 1 hpi and 3hpi (1 hpi image is shown). (A – B) Combined data from 3 independent experiments are shown. Per experiment >200 bacteria were scored in transfected, *i.e.* mCherry-positive cells. Error bars indicate SEM. Significance was determined by 2-way ANOVA. *** p<0.001; ****p<0.0001; n.s. = non-significant.

### Targeting of GBP1 to *S. flexneri* is dependent on its functional G domain, CaaX box and a C-terminal triple-arginine motif

To determine which protein motifs and properties of GBP1 (Figure 2A) render it uniquely capable to detect cytosolic Gram-negative bacteria, we generated and screened a large set of GBP1 mutant variants for co-localization with cytosolic *S. flexneri*. Mutant variants previously established to be defective for GTP hydrolysis (GBP1^R48A^) or nucleotide binding (GBP1^K51A^ and GBP1^S52N^) (21, 22) failed to associate with *S. flexneri* (Figure 2B). These findings were expected, because GTP binding and hydrolysis are required for GBP1 dimerization, protein polymerization and membrane binding (21–26). We also found that targeting to *S. flexneri* was dependent on the C-terminal CaaX box of GBP1 (Figure 2B). This finding was also expected as CaaX box-dependent prenylation of GBP1 provides a lipid anchor critical for the membrane association of GBP1 (25–27).

**Figure 2.**
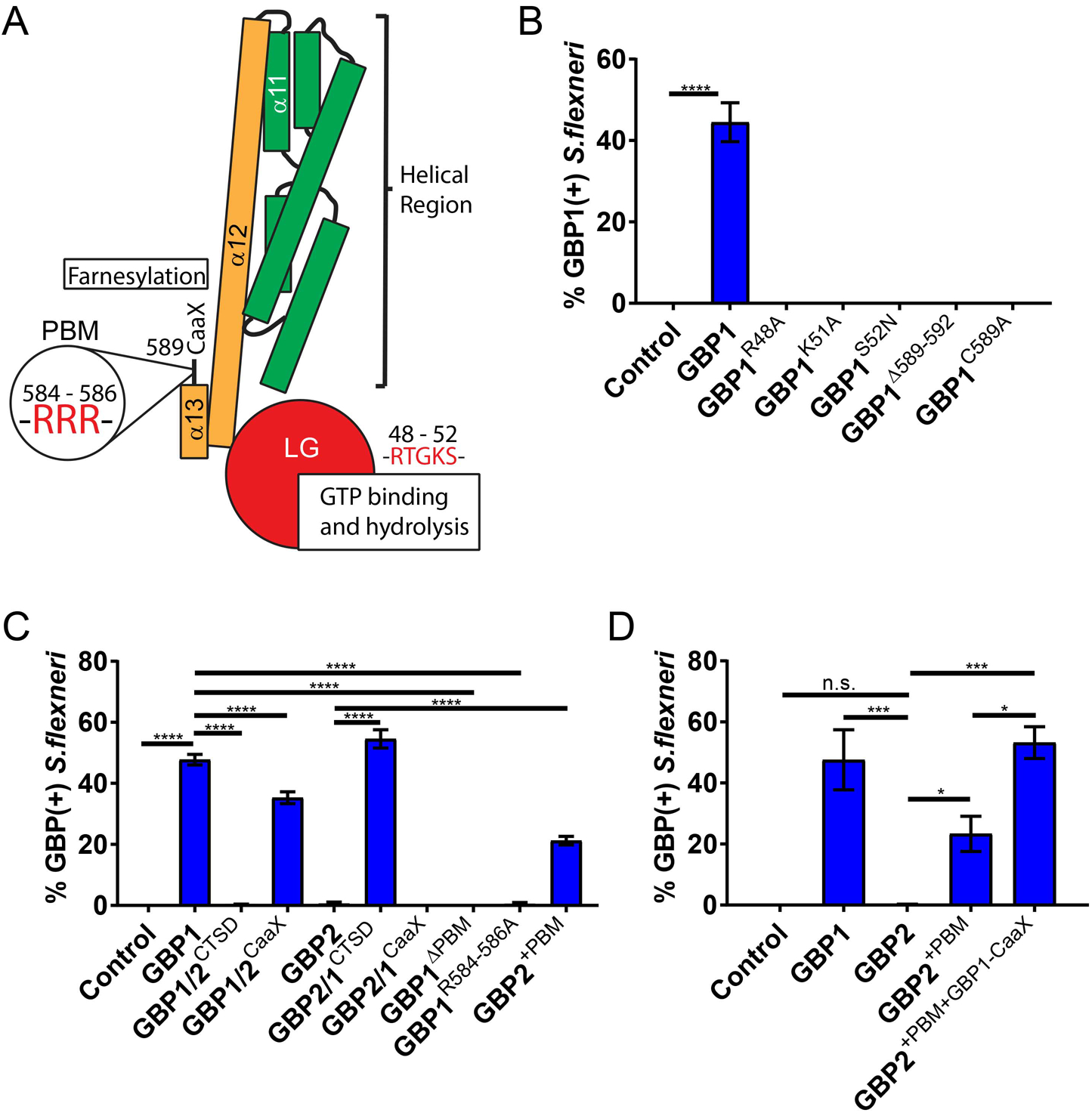
Targeting of GBP1 to *S. flexneri* is dependent on its functional G domain, CaaX box and a C-terminal triple-arginine motif. (A) Schematic depiction of critical protein motifs, domains and specific residues within the structure of human GBP1 is shown in. (B) A549 or (C – D) HEK 293T cells, respectively, were transfected with the indicated mCherry (control), mCherry-tagged wildtype and variant GBP1/GBP2 expression constructs and infected with GFP^+^ *S. flexneri* at an MOI of 50 before assessing co-localization at 3 hpi. Combined data from 3 independent experiments are shown. Per experiment >200 bacteria were scored. Error bars indicate SEM. Significance was determined by 1-way ANOVA relative to GBP1 (B) or as indicated (C and D). *p<0.05; *** p<0.001; ****p<0.0001; n.s. = non-significant.

Both GBP1 and GBP2 contain highly homologous N-terminal G domains and carry C-terminal CaaX boxes (11), yet only GBP1 efficiently associates with *S. flexneri* in A549 cells (Figure 1A). We therefore generated protein chimeras between GBP1 and GBP2 to map the motif that uniquely directs GBP1 to cytosolic bacteria. Most of the divergence between GBP1 and GBP2 sequences is found within the C-terminal subdomain (CTSD) consisting of α-helices α12 and α13, a short flexible region, and the CaaX box. We therefore swapped the CTSDs of GBP1 and GBP2 to generate two complementary chimeric proteins and found that the GBP1-but not GBP2-derived CTSD determined protein targeting to *S. flexneri* (Figure 2C). Swapping the CaaX boxes on the other hand only moderately reduced GBP1 targeting to bacteria (Figure 2C), indicating the existence of an additional motif within the GBP1-CTSD critical for bacterial recognition. Within the flexible region of CTSD we identified a GBP1-specific poly-basic protein motif (PBM), a 6 amino acid stretch containing 5 basic residues (KMRRRK). Deletion of these 6 residues or mutation of the triple-arginine cassette to triple-alanine abrogated co-localization of GBP1 with *S. flexneri* (Figure 2C). In a complementary approach, we found that insertion of the GBP1-PBM between GBP2 residues 586 and 587 (GBP2^+PBM^) was sufficient to drive significant targeting of GBP2 to *S. flexneri* (Figure 2C). Swapping the GBP2 with the GBP1 CaaX box further improved the bacteria-targeting efficiency such that GBP2^+PBM+GBP1-CaaX^ is recruited to *Shigella* at levels comparable to wildtype GBP1 (Figure 2D). These data demonstrate that a triple-arginine motif within the C-terminal PBM of GBP1 is essential and sufficient to equip both GBP1 and GBP2 with the ability to detect cytosolic *S. flexneri*, while the GBP1 CaaX box further improves targeting efficiency.

### Triple-arginine motif controls delivery of GBP1 to sterilely damaged vacuoles

To identify possible molecular targets for the C-terminal PBM of GBP1, we followed up on our previous published observation that GBP1 but none of its human paralogs detect sterilely damaged endogenous vacuoles in human embryonic kidney (HEK) 293T cells (28). The disruption of vacuoles leads to the cytosolic exposure of glycans that are normally confined to the vacuolar lumen. These exposed sugars then prompt the recruitment of the β-galactoside-binding lectin Galectin-3 as an established marker for loss of vacuolar integrity (29). To determine whether the PBM of GBP1 could promote the recognition of glycans, we tested its role in the delivery of GBP1 to ruptured vesicles. We first determined whether the structural requirements for the delivery of GBP1 to ruptured endosomes generally resembled those for the targeting of GBP1 to bacteria. In support of this premise, we observed that delivery of GBP1 to damaged endosomes was dependent on its functional G domain and CaaX box (Figure 3A). We further noticed that the CTSD of GBP1 was essential for the delivery of GBP1, and sufficient to promote recruitment of chimeric GBP2 protein, to damaged vesicles (Figure 3B). Deletion of PBM or mutation of the triple-arginine motif led to substantially reduced co-localization between Galectin-3-marked vacuoles and GBP1 (Figure 3B), indicating a functional role for the triple-arginine motif in the delivery of GBP1 to disrupted vesicles. These observations suggested that the triple-arginine motif of GBP1 directly or indirectly detects glycans.

**Figure 3.**
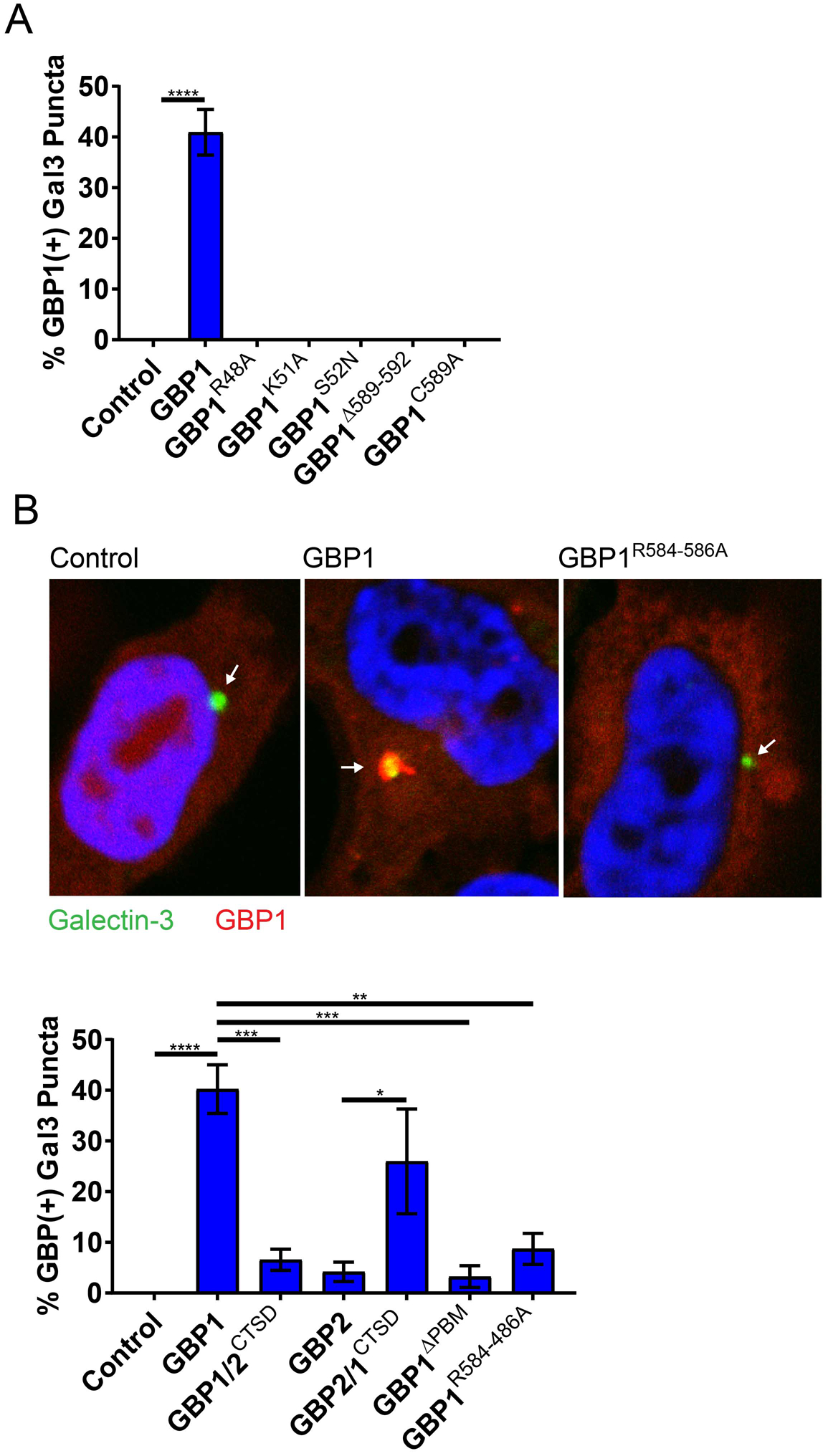
Localization of GBP1 to damaged endosomes is dependent on its triple-arginine motif. 293T cells stably expressing YFP-Galectin-3 (Gal3) were transiently transfected with the indicated expression constructs and approximately 1 day later treated with calcium phosphate precipitates to induce endosomal damage leading to YFP-Gal3 puncta formation. (A and B) Combined data from 3 independent experiments are shown. Per independent experiment >400 YFP-Gal3 puncta were scored for GBP1 co-localization in mCherry-positive cells. Error bars denote SEM. 1-way ANOVA was used to assess significance. **p<0.01; *** p<0.001; ****p<0.0001; n.s. = non-significant. (B) Arrows within representative images point to Galectin-3-positive vacuoles.

### *S. flexneri* mutants lacking O-antigen are targeted infrequently by GBP1

Because GBP1 detects broken vacuoles that are marked by cytosolically exposed polysaccharides normally confined to the vacuolar lumen, we speculated that bacterial surface-exposed polysaccharides could similarly underlie the recruitment of GBP1 to *S. flexneri.* We thus investigated whether GBP1 either directly or indirectly detects lipopolysaccharide (LPS), the main building block of Gram-negative bacterial envelope. Because O-antigen forms the outward sugar portion of LPS on the surface of bacteria, we tested GBP1 recruitment to the O-antigen-deficient *galU* and *rfaL* ‘rough’ *S. flexneri* mutants, which enter the host cell cytosol of non-polarized epithelial cells at comparable frequencies (30). We found that co-localization of ectopically expressed (Figure 4C) or endogenous GBP1 (Figure 4C) to *S. flexneri* rough mutants was substantially diminished relative to GBP1 targeting to wildtype *S. flexneri,* indicating that O-antigen recognition plays an important role in GBP1 recognition of cytosolic Gram-negative bacteria. We previously demonstrated that Galectin-3 interacts with a subset of murine GBPs and promotes their recruitment to *Legionella-*as well as *Yersinia*-containing vacuoles (28). Because Galectin-3 binds to O-antigen as well as the inner core of LPS (31), we hypothesized that Galectin-3 could promote the delivery of GBP1 to the Gram-negative bacterium *S. flexneri.* However, we observed that ectopically expressed GBP1 co-localized with *S. flexneri* with comparable frequencies in wildtype and Galectin-3 deficient mouse embryonic fibroblasts (MEFs) (Figure S2). Together, these data suggest GBP1 detects cytosolic *S. flexneri* in an O-antigen-dependent but Galectin-3-independent manner.

**Figure 4.**
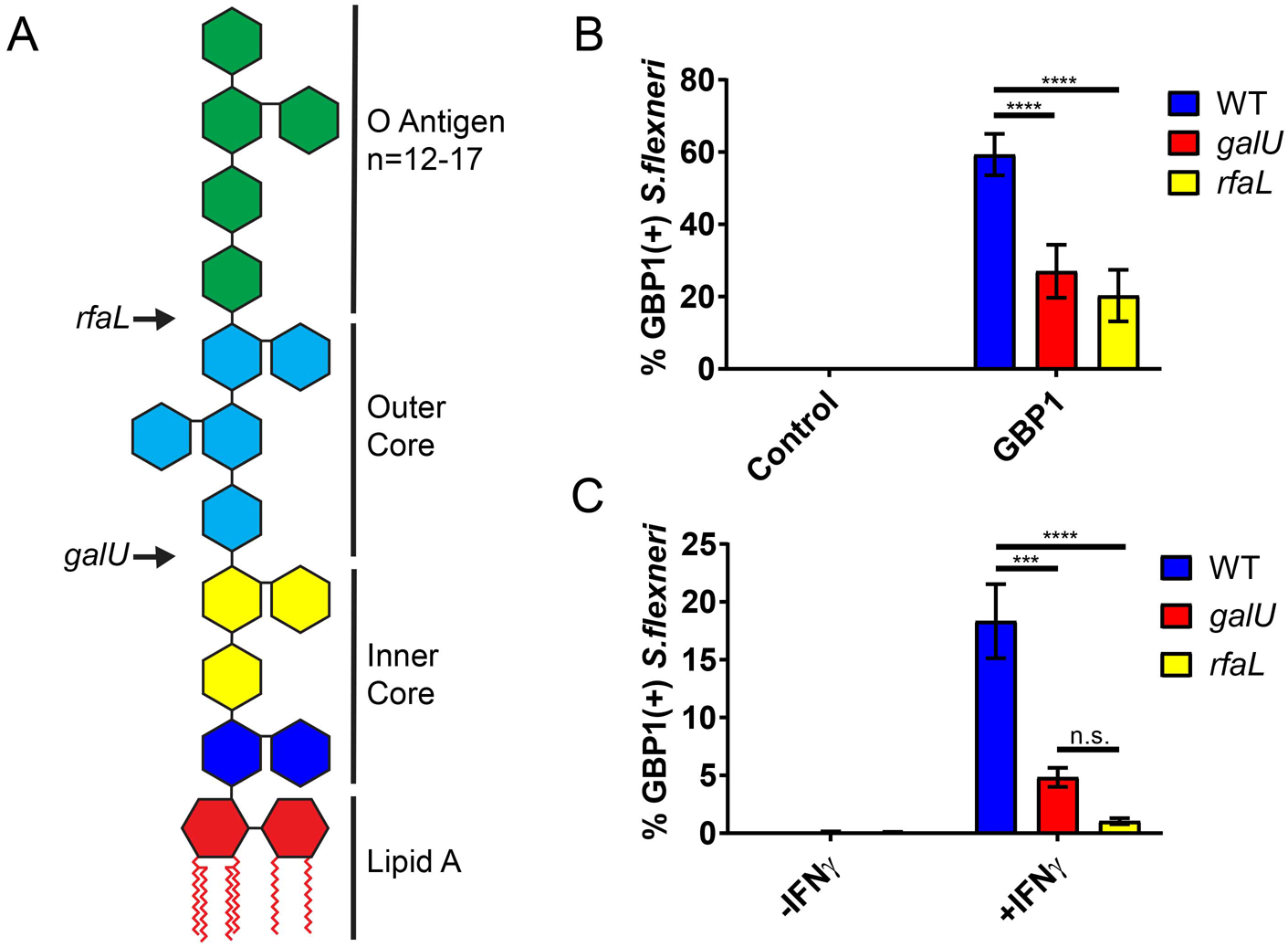
Bacterial targeting of GBP1 is diminished for *S. flexneri* rough mutants. (A) Arrows indicate site of truncation to LPS structure resulting from the loss-of-function mutations in bacterial genes *galU* and *rfa.* (B) 293T cells were transfected with mCherry-GBP1 or mCherry control, whereas (C) HeLa cells were left untransfected. Following infection with wildtype or the indicated *S. flexneri* mutants at an MOI of 50, co-localization of (B) ectopically expressed GBP1 or (C) endogenous GBP1 was quantified. Combined data from at least 3 independent experiments are shown. Per experiment >400 bacteria were scored per experiemental condition. Error bars represent SEM. 2-way ANOVA was used to assess statistical significance. *** p<0.001; ****p<0.0001; n.s. = non-significant.

### GBP1-decorated *S. flexneri* replicate within intracellular bacterial clusters

Previous work demonstrated the lytic destruction of GBP-bound cytosolic bacteria in mouse cells (13, 14). To assess whether recruitment of human GBP1 to *S. flexneri* would similarly drive bactericidal effects, we used an IPTG-inducible GFP reporter system as a bacterial viability indicator. In these experiments, we infected HeLa cells with reporter-positive *S. flexneri,* then induced GFP expression with IPTG at 2 hpi and stained cells for endogenous GBP1 and LPS at 4 hpi (Figure 5A). As expected, inhibition of bacterial translation with chloramphenicol treatment at 2 hpi blocked detectable GFP expression at 4hpi (Figure 5B), thus validating the reporter system. In the absence of chloramphenicol, GBP1-targeted and -untargeted bacteria were GFP-positive at comparable frequencies, indicating that GBP1 recruitment to *S. flexneri* fails to promote bacterial killing at this time point (Figure 5B). Instead, we noticed that GBP1-positive bacteria form intracellular clusters. To determine whether GBP1 could be responsible for this clustering effect, we generated HeLa cells that lack *GBP1* (*GBP1*^KO^) (Figure 5C). Following IFNγ priming to induce expression of GBPs, we observed reduced bacterial clustering in two independent *GBP1*^KO^ clonal cell lines as compared to the parental HeLa cells (Figure 5D). Bacterial clustering was restored in *GBP1*^KO^ cells by ectopically expressing wildtype GBP1 but not a triple-arginine mutant (Figure 5D). Furthermore, we found that GBP1-decorated, clustered bacteria not only remained viable but continued to divide (Figure 5E and video S2). Thus, our observations indicate that triple-arginine-dependent delivery of GBP1 to cytosolic *S. flexneri* promotes bacterial clustering but fails to execute bactericidal or bacteriostatic activities.

**Figure 5.**
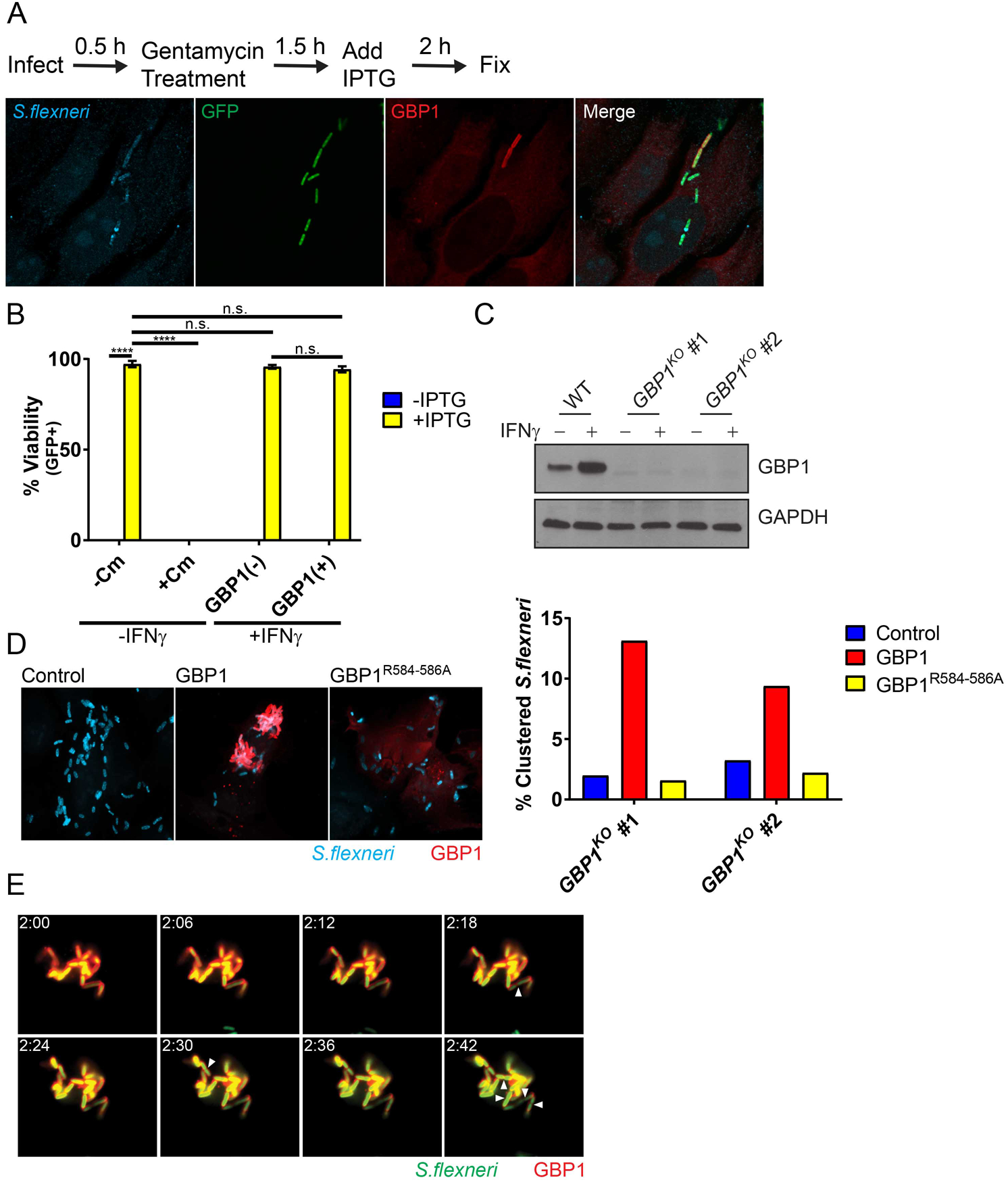
GBP1-tagged *S. flexneri* replicate within intracellular bacterial aggregates. (A) IFNγ-primed and unprimed HeLa were infected at an MOI of 50 with *S. flexneri* carrying an IPTG-inducible GFP reporter plasmid. IPTG, and where indicated also chloramphenicol (Cm), were added to the culture medium at 2 hpi. At 4 hpi cells were fixed and stained with anti-LPS (blue) and anti-GBP1 (red). A representative image of infected, IFNγ-primed HeLa cells is shown. (B) Combined data from 3 independent experiments as described under (A) are shown. (C) Protein lysates from IFNγ-primed parental HeLa (WT) and two independent *GBP1*^*KO*^ clones were transferred to membranes and probed with anti-GBP1 and anti-GAPDH antibodies. (D) Two independent *GBP1*^*KO*^ HeLa cell clones were transfected with wildtype GBP1 or GBP1^R584-586A^ mCherry fusion proteins and then infected with *S. flexneri* at an MOI of 50. Bacteria were visualized via with anti-LPS (blue) immunostaining. Cluster analysis was performed as described in Materials and Methods. (E) *GBP1*^*KO*^ HeLa cells were transfected with mCherry-GBP1 and infected with poly-D-Lysine pre-treated *S. flexneri* at an MOI of 10. Cells were infected for 30 minutes, washed and then placed in Phenol-red free DMEM. Video microscopy was begun at 2 hpi (2:00). Images of 6-minute intervals between 2:00 and 2:42 are shown. Arrowheads point at bacteria that have undergone cell division. 2-way ANOVA was used to assess statistical significance. ****p<0.0001; n.s. = non-significant.

### GBP1 inhibits actin-based motility in a triple-arginine motif-dependent manner

GBP1-decorated *S. flexneri* cluster intracellularly and therefore phenocopy *S. flexneri* Δ*icsA* mutants (Figure 6A) that lack the ability to form actin tails and hence are non-motile inside the host cell cytosol (32). Thus, we hypothesized that GBP1 blocks *S. flexneri* actin tail formation. In support of this hypothesis we observed a marked reduction in actin tails associated with GBP1-tagged versus -untagged *S. flexneri* in 293T cells expressing mCherry-GBP1 (Figure 6A). To complement these GBP1 overexpression studies, we scored actin tail-positive bacteria in wildtype versus *GBP1*^KO^ HeLa cells. We found that IFNγ priming reduced the number of bacteria with actin tails in wildtype but not in *GBP1*^KO^ cells (Figure 6B). Complementation of *GBP1*^KO^ cells with wildtype GBP1 but not the triple-arginine mutant (GBP1^R584-586A^) dramatically reduced the number of actin tail-positive bacteria to levels observed in IFNγ-primed wildtype cells (Figure 6B). As expected, GBP1-mediated inhibition of actin tail formation correlated with a GBP1-mediated block in cell-to-cell spread (Figure 6C). Inhibition of cell-to-cell spread was restored in *GBP1*^KO^ cells complemented with wildtype but not the triple-arginine mutant (GBP1^R584-586A^) (Figure 6D), indicating that GBP1 targeting to bacteria is required for inhibition of bacterial cell-to-cell spread.

**Figure 6.**
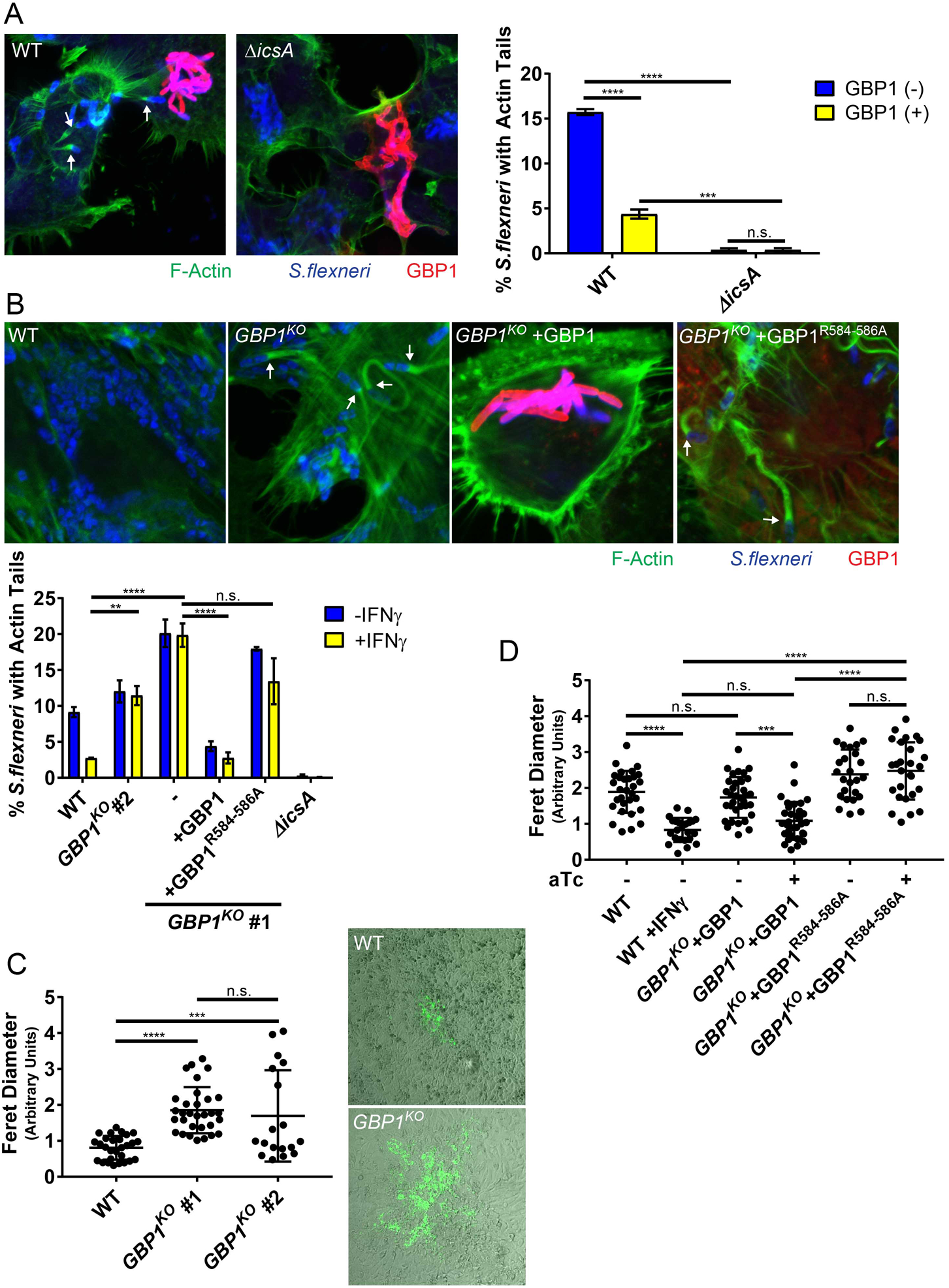
GBP1 restricts actin tail formation and cell-to-cell spread via its triple-arginine motif. (A) 293T cells were transfected with mCh-GBP1 and then infected with wildtype (WT) or Δ*icsA S. flexneri* at an MOI of 50. Confocal images were taken at 3 hpi and analysed for actin tail formation as described under Materials and Methods. Data are combined from 3 independent experiments. Per experiment and condition >550 bacteria were examined in mCherry-positive cells. (B) Parental HeLa-Cas9 (WT) and *GBP1*^*KO*^ cells were transfected with mCherry-GBP1 or mCherry-GBP1^R584-586A^ as indicated, then primed with IFNγ overnight and infected with *S. flexneri* at an MOI of 50. Cells were fixed and stained at 3 hpi and representative images are shown. Bacterial actin tail formation inside mCherry-positive cells was quantified and combined data from 3 independent data are shown. (C) Plaque assays on IFNγ-primed cells were performed as described under Materials and Methods and combined data from 3 independent experiments as well as representative images are shown. (D) HeLa-Cas9 (WT) cells or *GBP1*^*KO*^ (clone #1) with stably integrated pInducer-GBP1 or pInducer-GBP1^R584-586A^ expression constructs where either primed with IFNγ or stimulated with 1 µg/ml of aTc, as indicated. Combined data from 3 independent experiments are shown. Error bars depict SEM (A – C) or standard deviation (D). 2-way (A – B) or 1-way ANOVA (C – D) were used to determine statistical significance. **p<0.01; ***p<0.001; ****p<0.0001; n.s. = non-significant.

### Endogenous GBP1 associates with *S. flexneri* and recruits additional GBP paralogs in HeLa but not in A549 cells

GBP1 frequently forms heterodimers with other members of the GBP family such as GBP2 (3–5). To determine whether GBP1 recruits other GBP paralogs to cytosolic *S. flexneri,* we ectopically expressed mCherry-tagged human GBP1-7 in wildtype and *GBP1*^KO^ HeLa cells and monitored for subcellular localization of each with cytosolic *S. flexneri*. We found that ectopically expressed GBP2, and to a lesser extent GBP3, GBP4 and GBP6 co-localized with *S. flexneri* in HeLa cells in a GBP1-dependent manner (Figure 7A). This finding was curious, as we had not observed any co-localization of these GBP paralogs with *S. flexneri* in wildtype A549 cells (Figure 1A). We therefore monitored the subcellular localization of endogenous GBP1 in A549 cells and found that endogenous GBP1 targeted *S. flexneri* in HeLa but not in A549 cells (Figure 7B), which correlated with reduced expression of GBP1 protein in A549 compared to HeLa cells (Figure 7C). While endogenous GBP1 failed to decorate *S. flexneri* in A549 cells, we noticed that in the same cell line about 40% of cytosolic *B. thailandensis* stained positive for endogenous GBP1 (Figure 7B). These data suggested that *B. thailandensis* is more susceptible to GBP1 targeting than *S. flexneri.* To test this hypothesis, we generated a titratable, *i.e.* anhydrotetracycline (aTc)-inducible expression system for GBP1 (Figure 7D). As expected, higher GBP1 expression levels promoted more frequent co-localization with either bacterium. However, the overall frequency of GBP1 co-localization was significantly higher for *B. thailandensis* than for *S. flexneri* (Figure 7E). These data indicated that *S. flexneri* is more resistant to GBP1 targeting than *B. thailandensis* and therefore suggested that *S. flexneri* actively interferes with cytosolic detection by GBP1.

**Figure 7.**
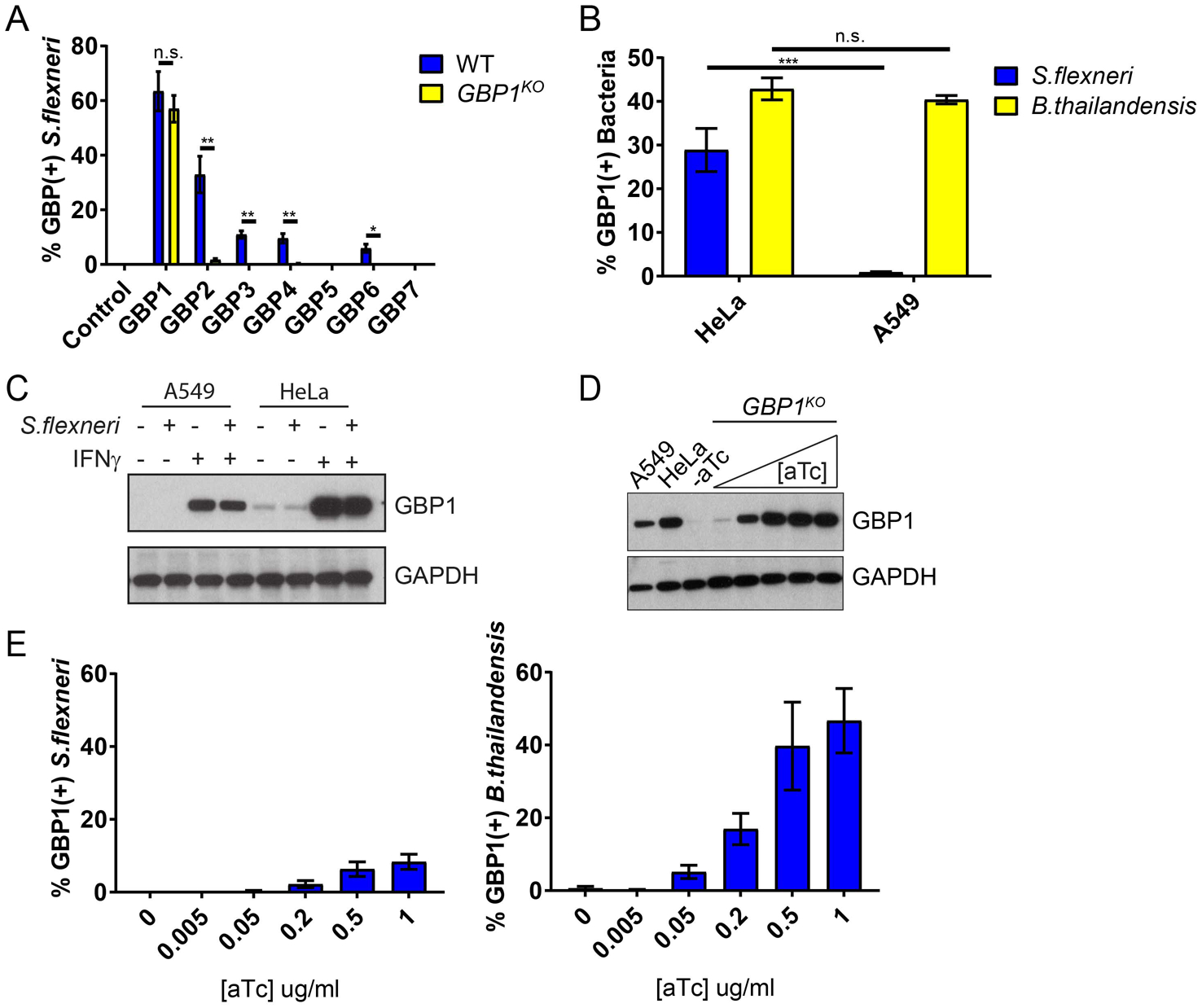
Endogenous GBP1 co-localizes with *S. flexneri* and recruits additional GBP paralogs in HeLa but not in A549 cells. (A) HeLa-Cas9 (WT) and *GBP1*^*KO*^ cells were transfected with indicated mCherry-GBP expression constructs and primed with IFNγ overnight. Cells were infected with GFP^+^ *S. flexneri* at an MOI of 50, fixed at 3 hpi and mCherry-positive cells were analysed for bacterial co-localization with GBPs. The combined data from 3 independent experiments are shown. Per experiment and condition >300 bacteria were scored. Error bars denote SEM and student t-tests were used to determine statistical significance. (B) IFNγ-primed HeLa and A549 cells were infected with GFP^+^ *S. flexneri* at an MOI of 50 or GFP^+^ *B. thailandensis* at an MOI of 100 and fixed and stained with anti-GBP1 at 3 hpi or 8 hpi, respectively. Combined data are from 3 independent experiments (>600 bacteria counted per experiment) are shown. Error bars represent SEM and 2-way ANOVA was used to assess significance. (C) Expression of endogenous GBP1 in IFNγ-primed HeLa and A549 cells was assessed by Western blotting. (D) *GBP1*^*KO*^ HeLa cells with stably integrated pInducer-GBP1 were stimulated overnight with indicated concentrations of aTc and protein lysates were subjected to Western blotting. Protein lysates from IFNγ-primed A549 and HeLa-Cas9 were included for comparison. (E) *GBP1*^*KO*^ HeLa cells with stably integrated pInducer-GBP1 were stimulated overnight with indicated concentrations of aTc, then infected with GFP^+^ *S. flexneri* (MOI of 50) for 3 h or GFP^+^ *B. thailandensis* (MOI 100) for 8 h and analyzed for bacterial co-localization with endogenous GBP1. Combined data from 3 independent experiments (>400 bacteria counted per experiment) are shown. Error bars represent SEM. *p<0.05; **p<0.01; ***p<0.001; n.s. = non-significant.

### IpaH9.8 blocks GBP1 recruitment and GBP1-mediated inhibition of actin-based motility

To account for our observations, we hypothesized that an effector secreted by the *Shigella* type 3 secretion system (T3SS) interferes with GBP1 targeting to cytosolic bacteria. To test this hypothesis, we monitored co-localization of GBP1 with two *S. flexneri* mutants deficient for the secretion of distinct subsets of type III effectors. A bacterial mutant lacking Spa15, the chaperone required for the secretion of the effectors IpaA, IpgB1, IpgB2, OspB, OspC1, OspC2, OspC3, OspD1 and OspD2 (33–36), was targeted with the same efficiency as wildtype *S. flexneri* (Figure 8A). However, Δ*mxiE Shigella,* a strain lacking the transcription factor that controls expression of second phase *S. flexneri* effectors including all IpaH effectors as well as OspB, OspC1, OspE1, OspE2, OspF, OspG and VirA (37–39), co-localized more frequently with GBP1 than wildtype *S. flexneri* in both A549 and HeLa cells (Figure 8A). These findings indicated that one or more mxiE-dependent T3SS effectors interfered with GBP1 function. To identify this virulence factor, we screened *S. flexneri* mutants deficient in individual mxiE-dependent T3SS effectors encoded on the *S. flexneri* virulence plasmid and found that Δ*ipaH9.8* mimicked the Δ*mxiE* phenotype (Figure 8B). Further, we found that Δ*ipaH9.8* was deficient for cell-to-cell spread in parental HeLa but not in *GBP1*^KO^ cells (Figure 8C), indicating that IpaH9.8-mediated interference with GBP1 targeting is critical for *S. flexneri* dissemination throughout the colonic epithelium. Cell-to-cell spread of the Δ*mxiE* mutant compared to wildtype *S. flexneri* was still moderately diminished in *GBP1*^KO^ cells, suggesting that one or more additional mxiE-controlled effectors other than ipaH9.8 promote cellular dissemination (Figure 8C).

**Figure 8.**
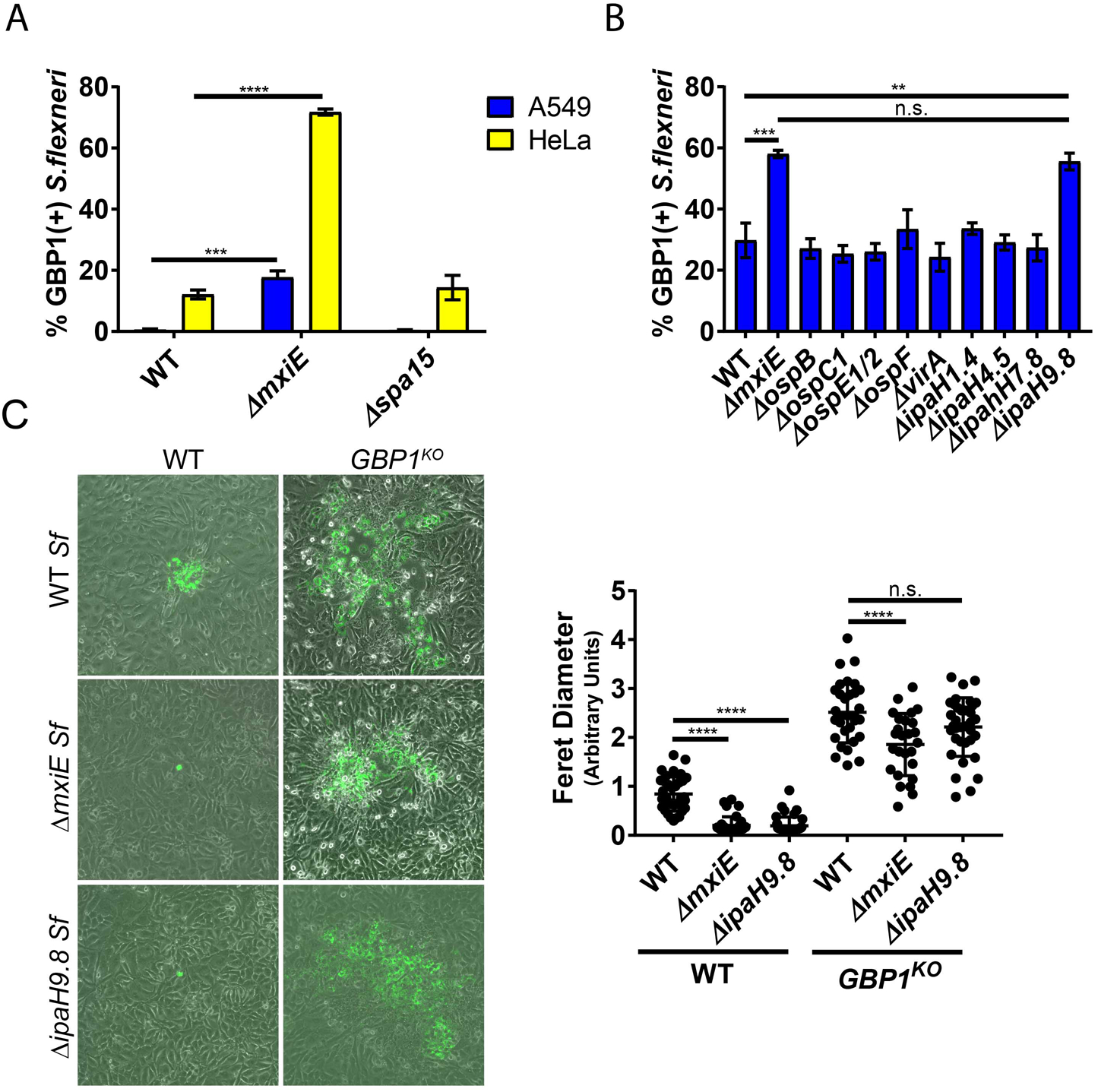
IpaH9.8 blocks GBP1 recruitment and GBP1-mediated inhibition of actin-based motility. (A) IFNγ-primed HeLa and A549 cells were infected with the indicated *S. flexneri* strains at an MOI of 50 and at 3 hpi processed and analyzed for co-localization with endogenous GBP1. Data are combined from 3 independent experiments scoring 600 bacteria per experiment. 1-way ANOVA was used to determine significance. (B) Experiments were conducted as in (A) with the *S. flexneri* mutant strains, as listed. 2-way ANOVA was used to determine significance of combined data from 3 independent experiments. (C) IFNγ-primed HeLa-Cas9 and derived *GBP1*^*KO*^ cells were infected with the indicated *S. flexneri* mutant strains. Representative images of plaques are shown. Combined data from 3 independent experiments are depicted. Error bars indicate SD. 2-way ANOVA was used to determine significance. **p<0.01; ***p<0.001; ****p<0.0001; n.s. = non-significant.

## Discussion

Many cell-autonomous defense mechanisms depend on the ability of the host cell to detect the precise location of an intracellular microbe inside an infected host cell (40). Pathogen-containing vacuoles are targeted by GBPs following immune recognition through missing-self or aberrant-self mechanisms (28, 41, 42), but the mechanism by which GBPs detect cytosolic pathogens had not previously been investigated. Here, we demonstrate that human GBP1 contains a unique triple-arginine motif within its flexible C-terminal region, which mediates the delivery of GBP1 to the Gram-negative cytosolic pathogen *S. flexneri.* We also demonstrate in this study that *S. flexneri* rough mutants lacking LPS O-antigen are targeted less efficiently by GBP1 than the co-isogenic wildtype strain. These observations suggest that the triple-arginine motif of human GBP1 either directly or indirectly interacts with the polysaccharide portion of LPS on the bacterial surface.

GBP1 binding to bacteria still occurs in the absence of O-antigen, albeit less efficiently. One possible explanation for this observation is that GBP1 associates with LPS through two or more distinct interactions, as has been described for Galectin-3. Galectin-3 binds to LPS through a high-affinity interaction with O-antigen as well as a low-affinity interaction with the inner LPS core (31). Alternatively, GBP1 could be recruited to bacteria through an additional LPS-independent mechanism. Future *in vitro* binding studies are required to test whether GBP1 binds to Gram-negative bacteria directly or indirectly, and to identify the specific binding substrate(s) or interaction partner(s) of the C-terminal triple-arginine motif essential for immune targeting of cytosolic *S. flexneri* by GBP1.

Previous work demonstrated that deposition of murine GBP2 on the cytosolic bacterial pathogen *Francisella novicida* results in bacteriolysis (13, 14). Similarly, murine GBPs also promote the lytic destruction of *S. flexneri* (43). Unexpectedly, we found that the delivery of human GBP1 to cytosolic *S. flexneri* is insufficient to kill bacteria or halt their replication. Instead, we observed that GBP1-bound bacteria fail to form actin tails and are consequentially restricted for efficient cell-to-cell spread. These observations were confirmed by an independent report published during the preparation of our manuscript (44). The differences between mouse and human in regards to the consequences of GBP targeting are intriguing and warrant future investigations to determine, for instance, whether the lytic pathway present in mouse cells is absent from human cells. The latter model is supported by a report demonstrating that GBP-dependent recruitment of the IFN-inducible GTPases Irgb10 to cytosolic *F. novicida* is required for bacteriolysis in mouse cells (45). Importantly, genes encoding Irgb10 and its paralogous subset of immunity related GTPases (IRGs) with a canonical GxxxxGKS P-loop sequence are absent from the human genome (46, 47), potentially accounting for the absence of an intracellular bacteriolytic pathway in human cells.

The dysregulation of cell-autonomous defense programs by *S. flexneri* is paramount to the microbes’ ability to colonize its human host and establish an infection (48). In agreement with two studies published during the preparation of our manuscript (43, 44), we found that the translocated bacterial ubiquitin E3 ligase IpaH9.8 inhibits the association of GBP1 with cytosolic *S. flexneri* in human cells. We did not investigate the mechanism by which IpaH9.8 interferes with GBP1 localization to bacteria, but the aforementioned studies demonstrated that IpaH9.8 ubiquitinates GBP1 and promotes its degradation via the proteasome (43, 44). While proteolytic degradation was clearly demonstrated at infections with high multiplicity of infection (MOI) or in cells ectopically expressing IpaH9.8 (43, 44), we failed to observe a pronounced decrease in GBP1 protein expression in cells infected with *S.flexneri* under the infection conditions used in our studies (Figure 7C). Furthermore, video microscopy data suggest that GBP1 ubiquitination by IpaH9.8 helps to dislodge GBP1 from targeted bacteria (44). We therefore propose that ubiquitination by IpaH9.8 not only tags GBP1 for proteolytic degradation but also directly blocks binding of GBP1 to bacteria and/or removes GBP1 from already targeted bacteria. Importantly, Li et al. demonstrated that IpaH9.8 promotes poly-ubiquitination of the GBP1 residues K582 and K587 (43), which flank the triple-arginine motif essential for recruiting GBP1 to *S. flexneri.* Whether PBM ubiquitination blocks GBP1 binding to bacteria through steric hindrance will be investigated in future studies.

## Materials and Methods

### Cell Lines and Culture

Primary WT and *Galectin-3*^*KO*^ (49) MEFs were made from C57BL/6J background mice acquired from the Jackson Laboratory. MEFs, A549, 293T, and HeLa cells were maintained at 37° and 5% CO_2_ in DMEM supplemented with 10% heat inactivated fetal bovine serum, nonessential amino acids (Gibco), and 55 μM β-mercaptoethanol. *GBP1*^*KO*^ lines were derived from HeLa-Cas9 cells, which contains a stably integrated Cas9 expression vector, lentiCas9-Blast (50). HeLa-Cas9 cells were transiently transfected with two guide RNAs directed against exon 6, GBP1-2 (TTGATCGGCCCGTTCACCGC) and GBP1-3 (TCCGGATACAGAGTCTGGGC), to introduce a predicted 64 bp deletion. Single clones were isolated by dilution. Resulting alleles are described in Table S1. All instances of IFNγ priming were done overnight at 200 U/ml.

### Bacterial Strains and Infections

Bacterial strains are summarized in Table S2. 2457T *S.flexneri*-derived *∆mxiE, ∆ospB, ∆ospC1, ∆virA, ∆IpaH1.4, ∆IpaH4.5, ∆IpaH7.8* and *∆IpaH9.8* were constructed using the λ red recombinase-mediated recombination system (51) using DNA oligomers listed in Table S3. GFP^+^ *S.flexneri* stains contain the vector pGFPmut2 (52). The IPTG-inducible GFP vector pRK2-IPTG-GFP was a gift from Wendy Picking. *S.flexneri* was grown at 37° on tryptic soy agar plates containing 0.01% Congo Red, as well as 50 µg/ml carbenicillin for pGFPmut2 and PRK2-IPTG-GFP containing strains. For infections, 5 ml tryptic soy broth (TSB) was inoculated with a single Congo red-positive colony and grown overnight at 37° with shaking. Saturated cultures were diluted 1:50 in 5 ml fresh TSB and incubated for 2.5-3 h at 37° with shaking. Bacteria were diluted in pre-warmed cell culture medium, and spun onto cells for 10 minutes at 700 x g. Plates were incubated at 37° for 30 minutes, then washed twice with Hanks Balanced Salt Solution (HBSS), followed by addition of cell culture medium containing 25 µg/ml gentamicin and further incubation at 37°. Unless otherwise specified, cells were infected at an MOI of 50. For time lapse microscopy and plaque assays, 5×10^7^ bacteria were harvested by centrifugation and incubated in 1 ml phosphate buffered saline (PBS) containing 5 μg/ml poly-D-lysine at 37° for 15 minutes with shaking, as described (53). Poly-D-lysine pre-treated bacteria were then used for infection, as above. *B. thailandensis* wiltype strain E264 carrying a GFP-expression construct was a gift from Edward Miao. *B.thailandensis +*GFP was grown at 37° on LB plates containing 100 ug/ml trimethoprim. For infections, overnight cultures were diluted 1:20 in LB + trimethoprim and grown for 3 h at 37° with shaking. Cells were infected at an MOI of 100, as above. Infected plates were incubated at 37° for 2.5 h, washed twice with HBSS, and incubated in the presence of 30 μg/ml kanamycin for 2 h. Kanamycin containing medium was then replaced with medium without antibiotics, and plates were incubated at 37° to 8 hpi. GFP^+^ wildtype *L. monocytogenes* strain 10403S was previously described (54) and grown at 37° on brain heart infusion (BHI) plates containing 10 µg/ml streptomycin and 7.5 µg/ml chloramphenicol. Infections were carried out in an identical manner to *S.flexneri* using saturated overnight cultures grown at 37° with shaking. Cells were infected with *L. monocytogenes* at an MOI of 5.

### Design of GBP Expression Constructs

Vectors containing mCherry-tagged constructs are derivatives of pmCherry-C1 (Clontech), and were constructed using DNA oligomers and restriction endonucleases listed in Table S4. For the chimeric constructs, GBP1/2^CTSD^ and GBP2/1^CTSD^, two synonymous mutations were introduced into GBP1 codons 478 and 479 of pmCherry-GBP1 using Quickchange site directed mutagenesis with GBP1-BclI oligomers (Table S5). These mutations introduced a BclI restriction endonuclease site within the sequence encoding the flexible region between helices 11 and 12. Following propagation of pmCherry-GBP1^BclI^ and pmCherry-GBP2 in *dam*^−^/*dcm*^−^ *E.coli* (New England Biolabs), BclI digestion and T4 Ligase-dependent repair was used to exchange the fragment encoding the C-terminal regions of the two proteins. All other mCherry-tagged chimeric and mutant constructs were constructed using Quickchange site directed mutagenesis with the primers listed in Table S5. GBP2^+PBM+GBP1-CaaX^ was constructed stepwise from pmCherry-GBP2^+PBM^ mutated with GBP2/1^CaaX^- and then GBP2^1587A^-oligomers. Tetracycline-inducible GBP1 was constructed by inserted the *GBP1* ORF into the third generation Tet-On vector, pInducer20 (55), using Gateway Cloning Technology with GBP1-attB oligomers listed in Table S5. GBP1^R584-586A^ was constructed by mutating the entry vector, pDONR221-GBP1, via Quickchange SDM using GBP1^R584-586A^-oligomers prior to Gateway LR reactions.

### Immunofluorescence Microscopy, actin tail analysis, Time-Lapse Microscopy and CPP assays

For standard microscopy cells were fixed in 4% paraformaldehyde and permeabilized with 0.25% Triton X-100 in PBS. Coverslips were probed with a 1:150 dilution of a rabbit monocolonal antibody against human GBP1 (Abcam ab131255) and/or a 1:50 dilution of a mouse monoclonal antibody against LPS (Raybiotech DS-MB-01267), followed by AlexaFluor conjugated secondary antibodies and 4 µg/ml of the nuclear dye Hoechst, where appropriate. For actin tail analysis, 1:40 Phalloidin conjugated to AlexFluor488 was added to the secondary antibody mix. For each field of view, Z-stacks were taken at 0.5 μm intervals on a Zeiss 710 inverted confocal to encompass the thickness of the cells. The percentage of bacteria with tails was determined by referencing across the collected Z-stacks through blinded analysis. For time-lapse microscopy *GBP1*^*KO*^ HeLa cells were plated in 35 mm glass bottom plates (MatTek) and transfected with pmCherry-GBP1. Cells were infected with GFP^+^ *S.flexneri* pre-treated with poly-D-lysine at an MOI of 10. Cells were imaged in DMEM without phenol red containing 5% FBS and 25 µg/ml gentamicin. Calcium phosphate precipitate (CPP) assays were carried out in 293T cells stably expressing YFP-Galectin-3 and transiently transfected with mCherry-tagged GBP constructs, as described in(28). Calcium phosphate precipitate (CPP) assays were carried out in 293T cells stably expressing YFP-Galectin-3 and transiently transfected with mCherry-tagged GBP constructs, as described in(28).

### IPTG-Inducible GFP Viability Assay

HeLa cells were stimulated overnight with 200 U/ml IFNγ and infected with *S.flexneri* containing the IPTG-inducible GFP vector, pRK2-IPTG-GFP. At 2 hpi, IPTG was added to a concentration of 1mM. Chloramphenicol was also added to a concentration of 60 µg/ml for negative controls. Coverslips were incubated for an additional 2 h before being fixed and stained with anti-LPS and anti-GBP1 antibodies, as described above.

### *S.flexneri* Cluster Analysis

Images were thresholded to remove background signal, and were subjected to connexity analysis to identify adjacent structures using the 3D objects counter in FIJI. Briefly, this process takes a pixel of interest and assesses whether the 26 surrounding pixels, (8 in plane, 9 above and below), contain signal above threshold. All adjacent pixels above threshold are then considered to be part of the same object. Each object is assigned a unique tag. The image is then subjected to a second pass to correctly identify connected items in three-dimensional space. The volume of each object is calculated and bacterial cluster analyzed. A volume representing individual and dividing bacteria was determined, and a threshold above this value was used to identify clusters.

### Plaque Assays

Plaque assays were performed essentially as previously described (56). Cells were plated to confluence in 35 mm glass bottom plates and stimulated overnight with 200 U/ml IFNγ. Cells were infected at an MOI of 5×10^−4^ with GFP^+^ *S.flexneri* pre-treated with poly-D-lysine. Following 30 minutes of infection at 37°, plates were washed twice with HBSS and overlaid with 1.5 ml DMEM without phenol red supplemented with 5% FBS and containing 25 ug/ml gentamicin, 200 U/ml IFNγ, 50 ug/ml carbenicillin, and 0.5% agarose. Plates were incubated for 10 minutes at room temperature before addition of 1 ml of cell culture medium without agarose. Plaques were visualized using phase contrast and fluorescence microscopy at 24 hpi, and the Feret diameter of individual plaques was determined using FIJI. For plaque assays using *GBP1*^KO^ cells complemented with pInducer-GBP1 or -GBP1^R584-586^, cells were stimulated overnight with 1 µg/ml aTc.

### Titration of GBP1 Expression from Tet-Inducible Expression System and Immunoblots

*GBP1*^*KO*^ cells stably transduced with pInducer-GBP1 constructs were incubated overnight with cell culture medium containing the indicated concentrations of aTc. Cells were lysed in RIPA buffer containing 1X protease inhibitor cocktail (Sigma) for 5 minutes on ice. Lysates were spun for 10 minutes at maximum speed in a microcentrifuge at 4° C, then combined with an equivalent volume of 2X Laemmli Sample Buffer containing β-mercaptoethanol, and incubated at 95° for 10 minutes. Samples were run on 4-15% Mini-PROTEAN TGX gels (Bio Rad) and transferred to nitrocellulose. Immunoblots were probed with 1:1,000 anti-GBP1 (Abcam ab131255), 1:1,000 anti-mCherry rabbit polyclonal (Abcam ab183628), or anti-GAPDH rabbit polyclonal (Abcam ab9485), followed by horseradish peroxidase-conjugated anti-rabbit IgG (Invitrogen).

### Statistical Analyses

Data analysis was performed using GraphPad Prism 6.0 software. Data shown are mean ± standard error of the mean (SEM) unless otherwise indicated. Statistical significance was calculated using 1-way or 2-way ANOVA or student t-test, as indicated. Significance was defined as (\*\*\*\**P* < 0.0001; \*\*\**P* < 0.001; \*\**P* < 0.01; \**P* < 0.05).

## Author contribution

Conceptualization: A.S.P. and J.C.; Methodology: A.S.P. and J.C.; Investigation: A.S.P., D.H., S.L., E.M.F., R.S., S.W.; Writing– Original Draft: J.C.; Writing – Review & Editing: A.S.P. and J.C.; Funding Acquisition: C.F.L. and J.C.; Resources: E-M.F., C.F.L.; Supervision: J.C.

## Acknowledgement

We thank Edward Miao from UNC Chapel Hill for sharing *B. thailandensis* strains, Marcia Goldberg from Harvard Medical School for providing the Δ*iscA S. flexneri* mutant and co-isogenic wildtype strains, Anthony Maurelli from the University of Florida for providing the *S. flexneri galU* and *rfaL* mutant and co-isogenic wildtype strains and Wendy Picking for providing pRK2-IPTG-GFP. We thank So Young Kim from the Duke Functional Genomics core for the production of HeLa *GBP1*^*KO*^ cells. We thank members of the Coers and Tobin labs for helpful discussions.

## Supplementary text

**Figure S1.**
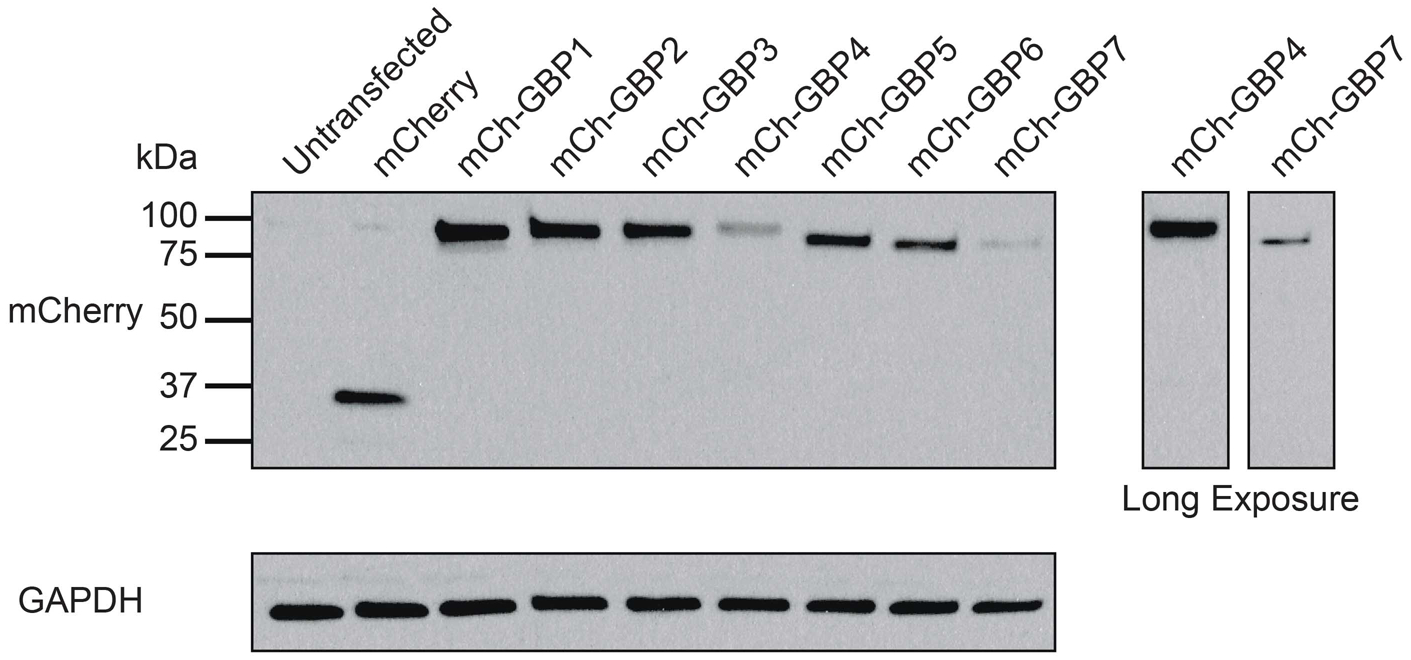
Expression of N-terminal mCherry-GBP fusion proteins in 293T cells. 293T cells were transiently transfected with mCherry-tagged human GBPs. Cells were lysed in RIPA buffer approximately 24 h post transfection and probed with anti-GBP1 and anti-GAPDH antibodies.

**Figure S2.**
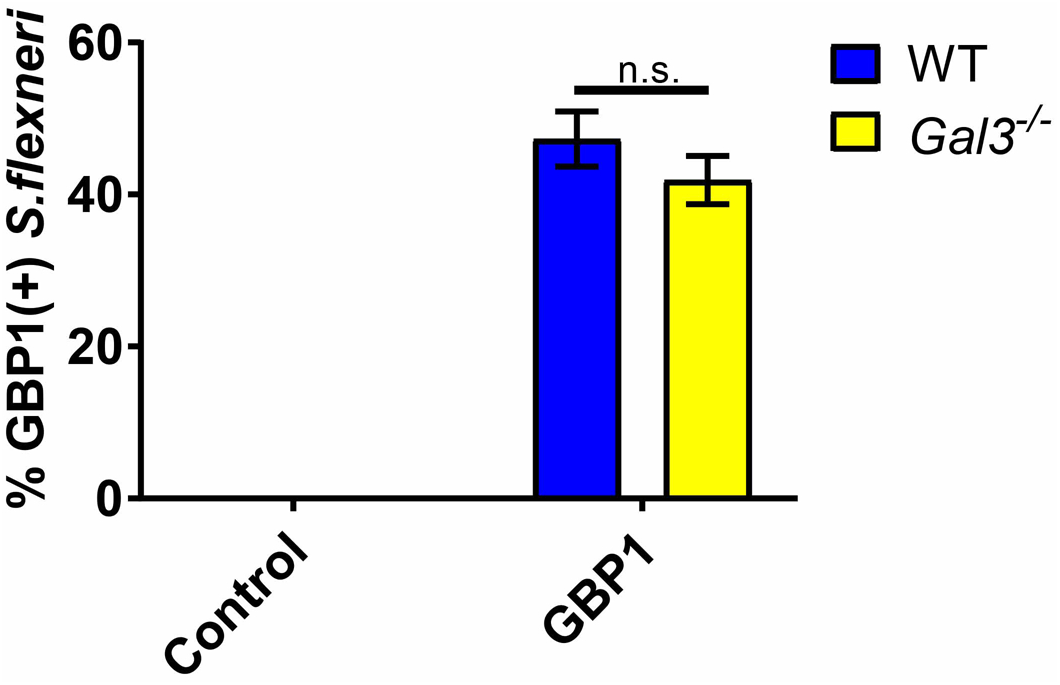
GBP1 co-localizes with *S. flexneri* independent of Galectin-3. Primary WT (C57BL/6J) and *Galectin-3*^−/−^MEFs were transiently transfected with expression constructs for mCh-GBP1 or an mCherry control, and subsequently infected with GFP^+^ *S. flexneri* at an MOI of 50. Cells were fixed at 3 hpi, and the percentage of GBP1-positive bacteria was determined by microscopy. Graph represents the mean percentage of GBP1-positive bacteria from three independent experiments. Error bars denote SEM. A 2-way ANOVA was used for statistical analysis. n.s. = non-significant.

### Supplementary Videos

**Video S1.** *GBP1*^*KO*^ HeLa cells transfected with mCherry-GBP1 were infected with poly-D-lysine pre-treated *S.flexneri* expressing GFP at an MOI of 10. Images were collected every 30 seconds for 3 hours, beginning at 10 minutes post infection.

**Video S2.** *GBP1*^*KO*^ HeLa cells transfected with mCherry-GBP1 were infected with poly-D-lysine pre-treated *S.flexneri* expressing GFP at an MOI of 10. Images were collected every 90 seconds for 45 minutes, beginning at 190 minutes post infection.

**Table S1.** GBP1 alleles in HeLa GBP1^KO^ clones #1 and #2. Deletion is annotated based on the corresponding nucleotide positions in GBP1 ORF. The predicted protein size of truncated protein resulting from the mutation is provided (full-length GBP1 is 592 amino acids in length).

| Mutation | Notes | Protein Size |
| --- | --- | --- |
| deletion 668-729 | Deletion resulting in frame shift and early stop codon | 234 aa |
| deletion 667-730 | Deletion resulting in frame shift and early stop codon | 245 aa |

**Table S2.**
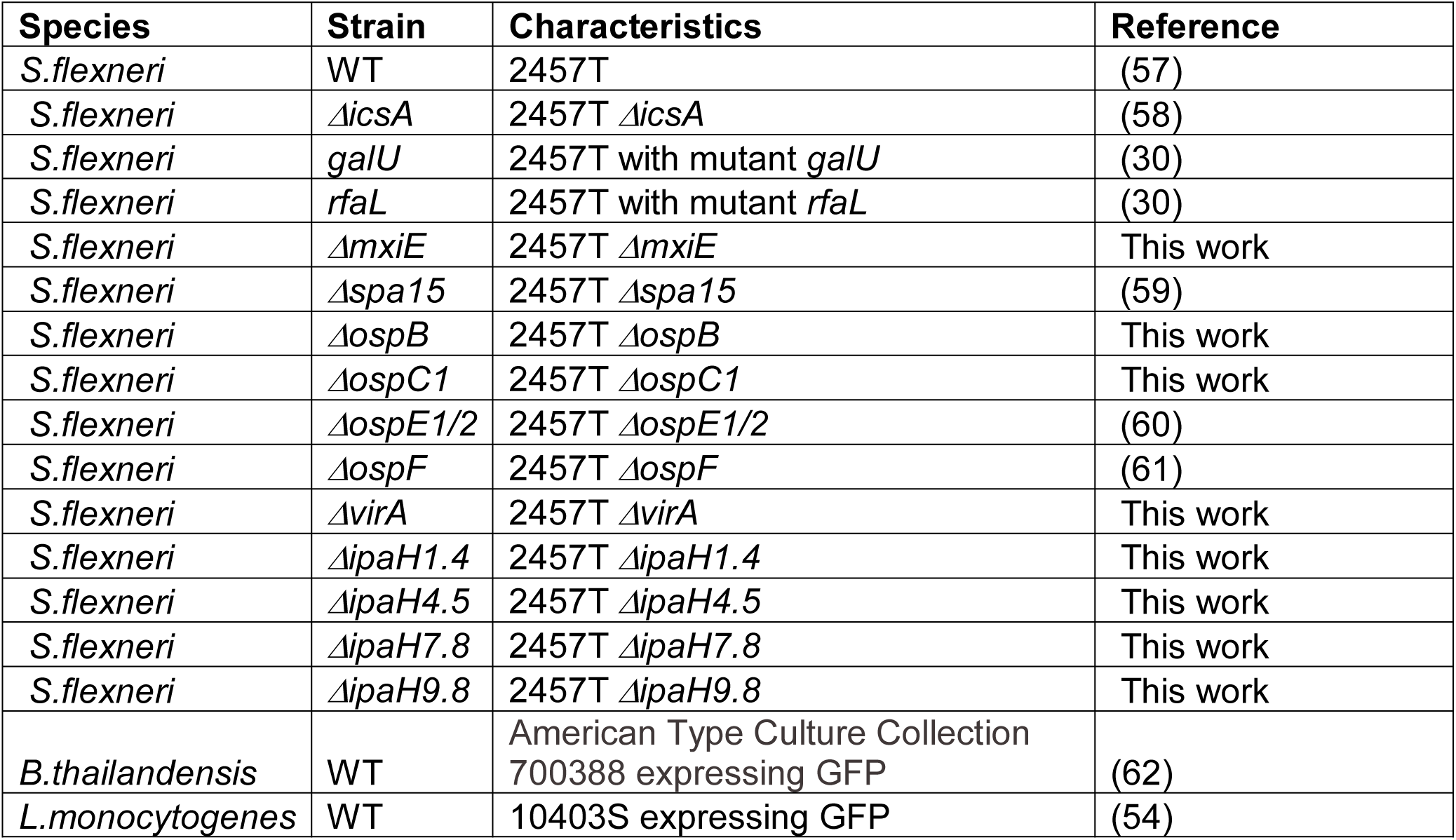
List of bacterial strains used in this study

**Table S3.**
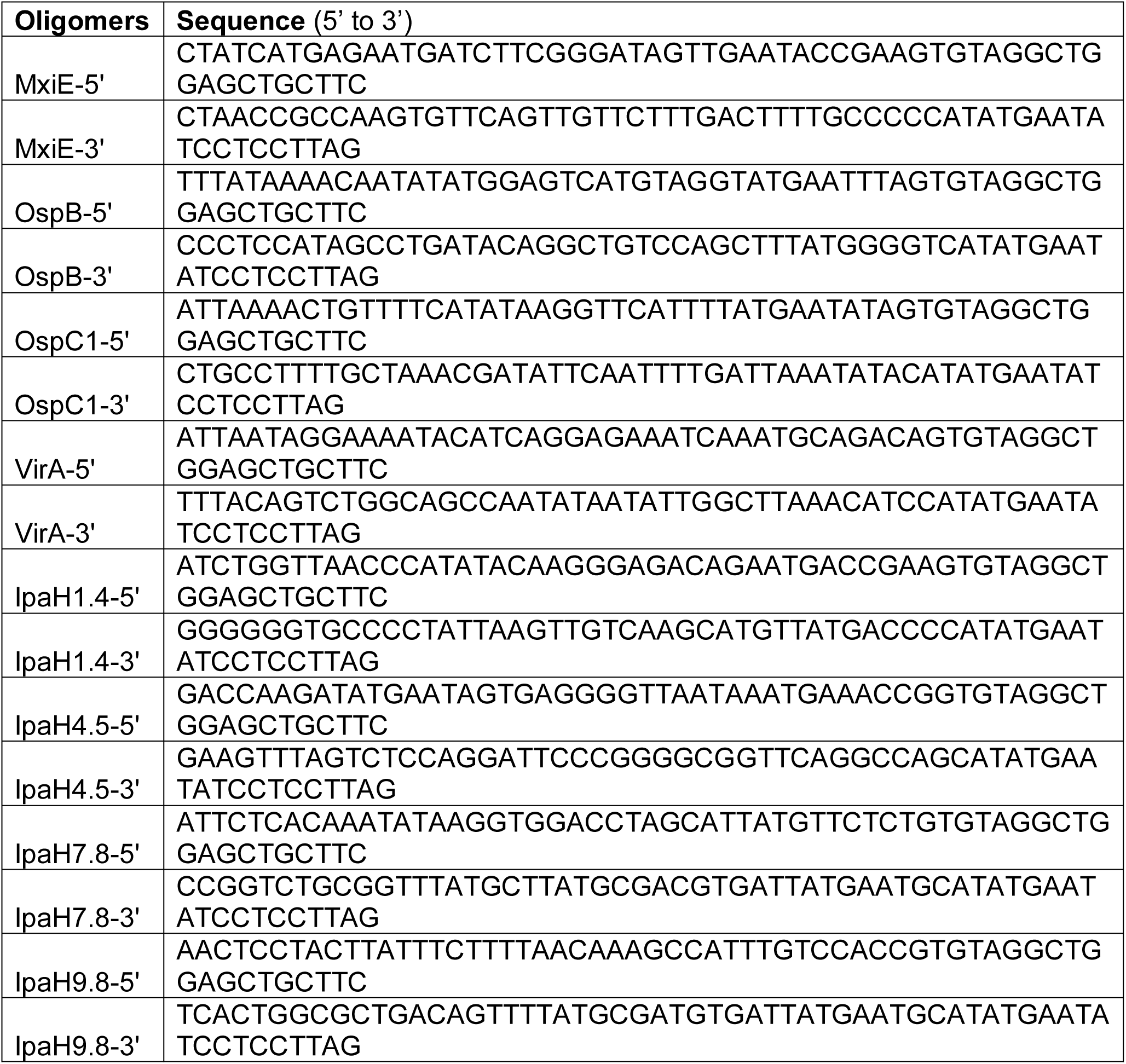
List of DNA oligomers used to generate *S. flexneri* mutant strains

**Table S4.**
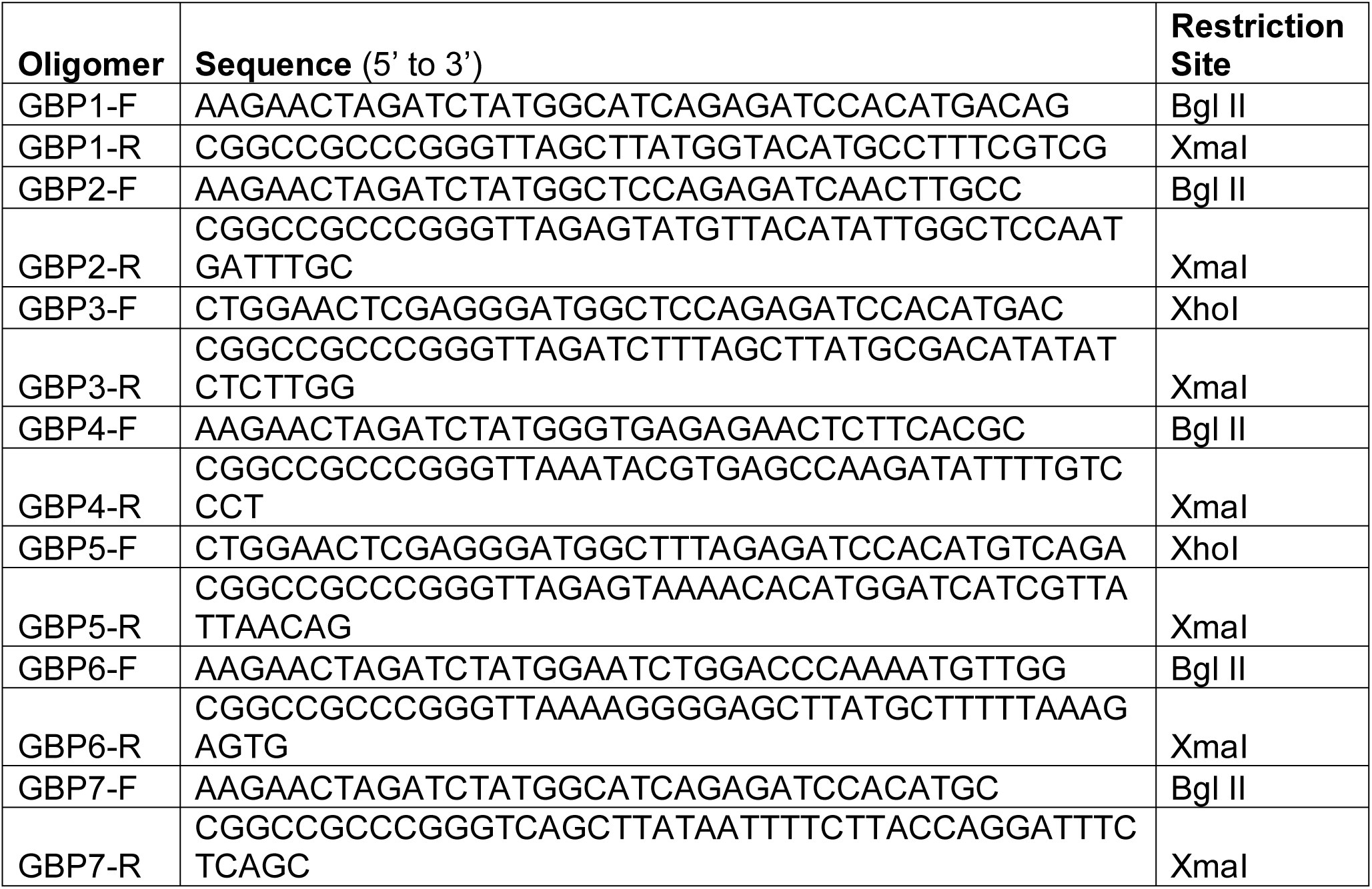
List of oligomers and restriction sites used to generate mCherry GBP fusion expression constructs

**Table S5.**
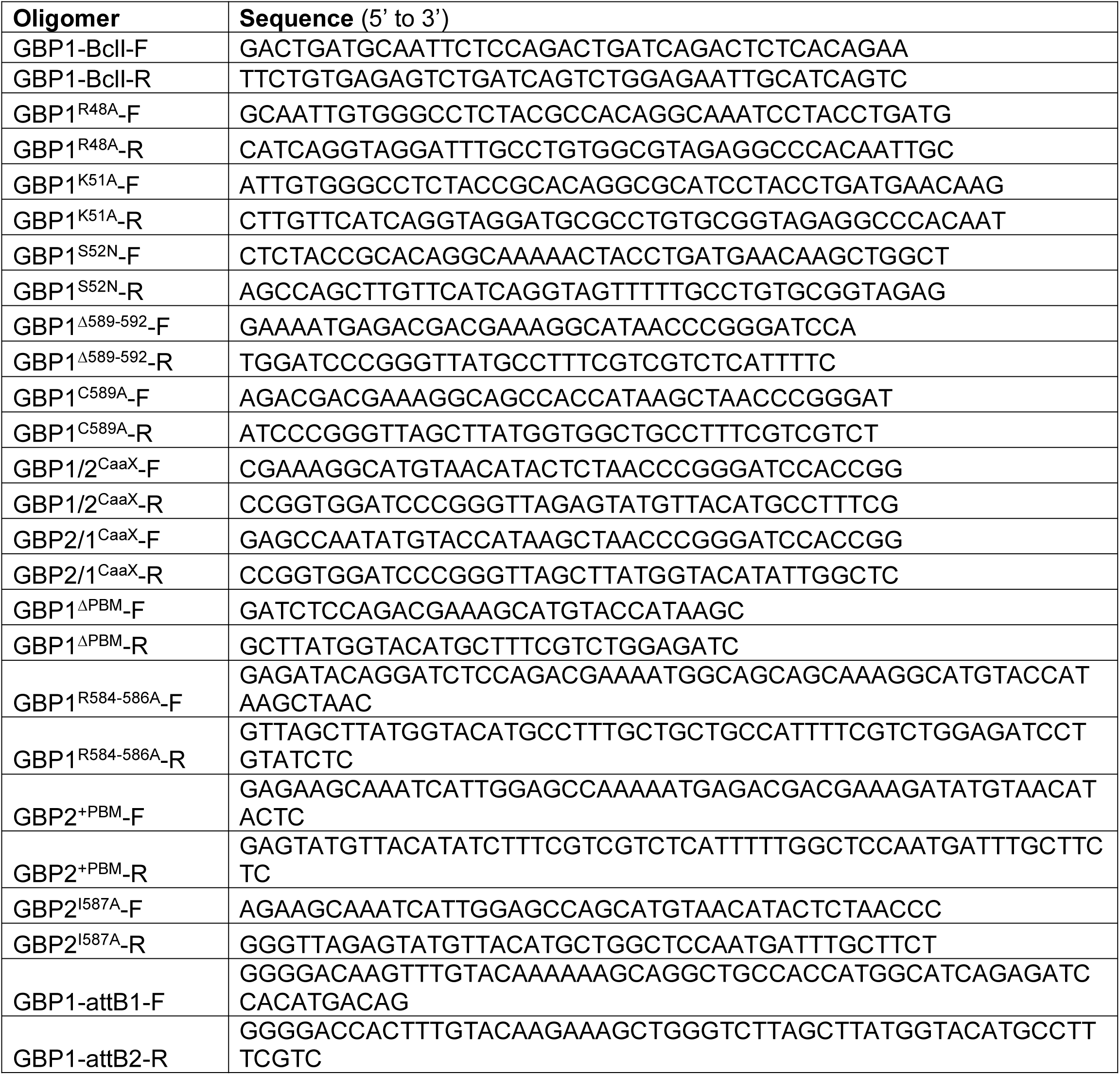
Oligomers used to generate GBP mutant and chimeric variants.

